# A general theory for temperature-dependence in biology

**DOI:** 10.1101/2021.04.26.441387

**Authors:** José Ignacio Arroyo, Beatriz Díez, Christopher P. Kempes, Geoffrey B. West, Pablo A. Marquet

## Abstract

At present, there is no simple, complete, and first principles-based model for quantitatively describing the full range of observed biological temperature responses. Here, we derive a theory exhibiting these features based on the Eyring-Evans-Polanyi theory governing chemical reaction rates, and which is applicable across all scales from the micro to the macro. Assuming only that the conformational entropy of molecules changes with temperature, we derive a theory for the temperature dependence which takes the form of an exponential function modified by a power-law. Our framework leads to six deductions applicable to any biological trait that depends on temperature, and elucidates novel aspects of universal temperature responses across the tree of life, from quantum to classical scales. All predictions are well supported by data for a wide variety of biological rates and steady states, from molecular to ecological scales and across multiple taxonomic groups. In addition, we provide novel explanations of several empirical relationships including optimal values in temperature response curves.

**One-Sentence Summary:** We derive a simple and universal formulae to characterize temperature responses of biological processes across the tree of life.

## Introduction

### Temperature dependence models and the Eyring-Evans-Polanyi (EEP) theory

Temperature is a major determinant of reaction rates of enzymes, which regulate processes that manifest at all levels of biological organization from molecules to ecosystems [1-7]. Formulating a fundamental theory for the response of biological rates to changes in temperature, especially in ecological systems, has become a matter of some urgency with the intensification of the climate crisis, particularly since existing models are unable to account for such responses across the entire range of temperatures that support life. Here we address this challenge by developing a comprehensive theory that unifies several key properties that have not been simultaneously included in past work. This is critical for making accurate predictions of biological quantities that are relevant in industrial applications, food production, disease spread, and responses to climate warming, among others. The model we derive is: i) based on first principles and fundamental chemical mechanisms; ii) mathematically simple in form, yet efficient in that it generates many predictions with very few free parameters; iii) general and applicable across multiple levels of biological organization and taxa, thereby manifesting a universal biophysical law. Among its many novel results, our theory makes six significant categories of new deductions that are confirmed by data and resolves unexplained observations in the temperature response of organisms.

Different models have been suggested to explain temperature dependence in biology, among which the Arrhenius equation [8-9] has become the most used by biologists and ecologists, as epitomized, for example, by the Metabolic Theory of Ecology (MTE), [7] and is given by

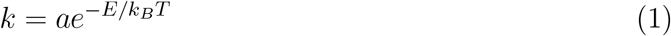

where *k* is some biological quantity (e.g. at the molecular level, enzyme reaction rate), *k*_*B*_ is Boltzmann’s constant, *T* is absolute temperature in Kelvin degrees (K), *E* is an effective activation energy for the process of interest, and *a* is an overall normalization constant characteristic of the process. Consequently, a plot of log *k* vs. 1*/T* should yield a straight line, often referred to as an Arrhenius plot. This equation was originally an empirical formulation, but was later motivated heuristically from chemical reaction theory [10, 11]. Although it has been instrumental in explaining the approximately universal temperature dependence across many diverse biological rates [5, 7], it cannot account for deviations that occur beyond certain temperature ranges in, for example, the metabolic rates of endotherms, thermophiles and hyperthermophiles [3, 5, 12]. Furthermore, experiments and observations have long established that the form of the temperature response has an asymmetric concave upward or downward pattern relative to the canonical straight-line Arrhenius plot. Consequently, there are ranges of temperatures where the traditional Arrhenius expression, Eq. (1), even gives the wrong sign for the observed changes in biological rates: they *decrease* with increasing temperature rather than increase, as predicted by Eq. (1).

The EEP transition state theory (TST) [13-14], which is the widely accepted theory of enzyme chemical kinetics, offers the possibility of developing a fundamental theory for the temperature dependence of biological processes that extends and generalises the heuristic Arrhenius equation by grounding it in the underlying principles of thermodynamics, kinetic theory and statistical physics [15]. The framework of the TST conceives of a chemical reaction as a flux of molecules with a distribution of energies and a partition function given by the Planck distribution, flowing through a potential energy surface (PES) which effectively simulates molecular interactions. The configuration of molecules flowing through this surface proceeds from i) a separate metabolite and enzyme to ii) an unstable metabolite-enzyme complex, which, iii) after crossing a critical energy threshold barrier, or transition state, then forms the final product (the transformed metabolite). EEP thereby derived the following equation for the reaction rate [11]

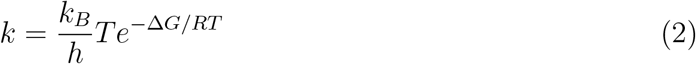

where *h* is Planck’s constant, Δ*G* is the change in Gibbs free energy or free enthalpy, *R* = *Nk*_*B*_ is the universal gas constant and *N* is Avogadro’s number. An overall coefficient of transmission also is originally part of (2) but is usually taken to be 1. The change in Gibbs free energy is the energy (heat) transferred from the environment to do chemical work. It can be expressed in terms of enthalpy (Δ*H*) and the temperature-dependent change in entropy, or dissipated energy (Δ*S*) [16], as Δ*G* = Δ*H* − *T* Δ*S*. Eq. (2) can then be written as:

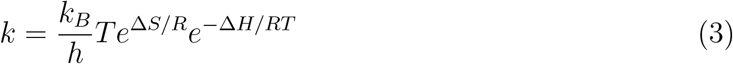

Analogous to the Arrhenius expression, Eqs. (2) and (3) describe an exponential response of the rate *k* to temperature provided, however, that there is no temperature dependence of the thermodynamic parameters. Models have been developed for including this temperature dependence, but they typically invoke several additional assumptions and new parameters [11, 17-18]. Furthermore, unlike the widespread use of the Arrhenius equation in the MTE, most models for temperature response have been conceived for a single level of biological organization (primarily at the enzymatic/molecular level) [6, 18] or for specific taxonomic groups; e.g. only for mesophilic ectotherms [19], endotherms [20], or thermophiles [21].

### Derivation of the Theory

Temperature changes the conformational entropy of proteins [23], which in turn determines the binding affinity of enzymes [24-25] and affects the flexibility/rigidity and stability of the activated enzyme-substrate complex and hence the reaction rate [25]. The resulting temperature dependence of the change in entropy, Δ*S* (with enthalpy and heat capacity remaining constant), is the simplest mechanism for giving rise to curvature in an Arrhenius plot and naturally leads, via Eq. (3), to power law deviations from the simple exponential form [22]. Following [16], the change of entropy for a given change in temperature can be expressed as *Td*Δ*S/dT* = Δ*C*, where Δ*C* is the heat capacity of proteins. Integrating over temperature gives Δ*S* = Δ*S*_0_ + Δ*C* ln (*T/T*_0_), where Δ*S*_0_ is the entropy when *T* = *T*_0_, an arbitrary reference temperature, commonly taken to be 298.15 *K* (25°*C*) [11]. Using this expression for Δ*S* in eq. (3), and after simplifying, we straightforwardly obtain [11]

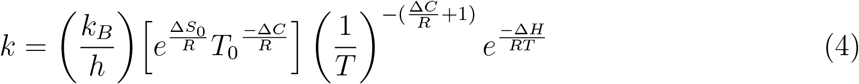

Eq. (4) has the form of a classic Arrhenius-like exponential term, modified by a powerlaw, but with a different interpretation of the “effective activation energy” in terms of the change in enthalpy. The pattern described by Eq. (4) is a curved temperature response in an Arrhenius plot of ln *k* vs. *T* ^−1^:

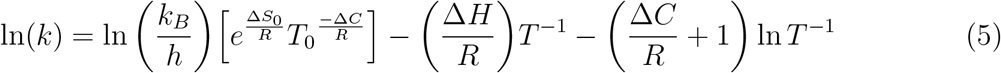

Consequently, *d* ln(*k*)*/dT* ^−1^ = −Δ*H/R* − (Δ*C/R* + 1)*/T* ^−1^, leading to the extrema of ln(*k*) occurring at 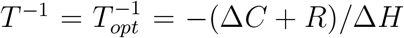 (see Supplementary Text S6). This is a minimum, i.e., the curve is concave upwards, or a “happy mouth”, if Δ*C >* −*R*, whereas it is a maximum, or a convex downwards “sad mouth”, if Δ*C <* −*R*. Furthermore, for 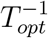 to be positive this requires Δ*H <* 0 for a minimum or Δ*H >* 0 for a maximum.

Several important points should be noted about our result:

1. Its simple mathematical form, namely an exponential modified by a power law, coincides with an empirical phenomenological equation suggested by Kooij in 1893 [26]. However, our derivation provides an underlying mechanism for the origin of the expression and, consequently, for how its parameters are related to the thermodynamic variables. Our approach differs from previous expressions derived from considerations of chemical kinetics [11]. For instance, a heuristic derivation inspired by a Maxwell-Boltzmann distribution predicts a similar expression but with a power law modification of *T* ^1*/*2^ rather than 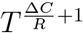 [13, 14], which apart from not having a mechanistic basis, is also unable to explain concave deviations.
2. An important consequence of our derivation is that it shows that a change of entropy with temperature is both sufficient and necessary for simultaneously explaining both the convex and concave curvatures commonly observed in temperature-response plots. Under a thermodynamic interpretation, the decrease in enzymatic rate with increasing entropy due to increasing temperature beyond the optimal, means that the disorder of the enzyme, and particularly of the active site, has reached a state that causes a decrease in the binding affinity to the ligands. In contrast, changes in enthalpy alone can only explain convex curvature but not concave. To see this explicitly, we express Δ*H* in terms of heat capacity in eq. (3), Δ*H* = Δ*H*_0_ − Δ*C*(*T* − *T*_0_), to obtain 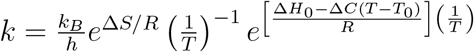, which leads to 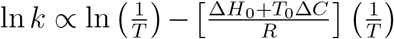. Regardless of the sign of both Δ*C* and/or Δ*H*_*0*_, this always results in a convex downwards curve and so cannot explain, nor accommodate, concavity. Hobbs et al. [27] included changes in both enthalpy and entropy with temperature and derived a significantly more complicated expression than ours based on TST. In contrast, the minimalist scenario developed here is one in which only changes in entropy with temperature need be considered.
3. The above derivation was for reaction rates at the microscopic enzymatic scale. Following the argument in the MTE we now show how it can be extended to biological variables at multiple scales up through multicellular organisms to ecosystems. The most salient example is metabolic rate, *B*. In general, this is derived by appropriately summing and averaging over all enzymatic reaction rates contributing to metabolism - some connected in series, some in parallel - and then summing and averaging over all cells: symbolically, 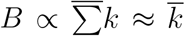. Assuming there is a dominant set of rate limiting reactions contributing to the production of ATP [19], then the temperature dependence of 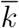, and therefore *B*, can be approximated by an equation of the form of Eq. (4), but with the parameters being interpreted as corresponding averages, 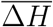 and 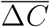. This results in: 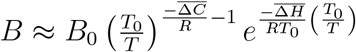, where *B*_0_ is a normalization constant (see Supplementary Material Eq. (S3.3) and Text S8).
4. Care, however, has to be taken with the normalization constants, such as *B*_0_ in the case of metabolic rate, since from Eq. (4), these would naively be proportional to the ratio of the two fundamental constants, *k*_*B*_ and *h*. The presence of Planck’s constant, *h*, for microscopic enzymatic reactions appropriately reflects the essential role of quantum mechanics in molecular dynamics. On the other hand, for macroscopic processes, such as whole body metabolic rate, the averaging and summing over macroscopic spatio-temporal scales which are much larger than microscopic molecular scales must lead to a classical description decoupled from the underlying quantum mechanics and, therefore, must be independent of *h*. This is analogous to the way that the motion of macroscopic objects, such as animals or planets, are determined by Newton’s laws and not by quantum mechanics, and therefore do not involve *h*. Formally, the macroscopic classical limit is, in fact, realised when *h* → 0. The situation here is resolved by recognising that the partition function for the distribution of energies in the transition state of the reaction has not been explicitly included in Eq. (2). This is given by a Planck distribution which leads to an additional factor 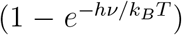. where *ν* is the vibrational frequency of the bond, as first pointed out by Herzfeld [28]. For purely enzymatic reactions discussed above this has no significant effect since *k*_*B*_*T << hν*, and thus 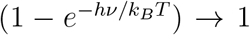, resulting in Eq. (2). Multicellular organisms, however, correspond to the classical limit where *h* → 0 so *k*_*B*_*T >> hν* and 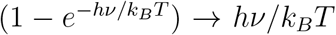, thereby cancelling the *h* in the denominator of Eq. (4). Consequently, the resulting temperature dependence of macroscopic processes, such as metabolic rate, become independent of *h*, as they must, but lose a factor of *T* relative to the microscopic result, Eq. (4), so for metabolic rate, *B*, this is:

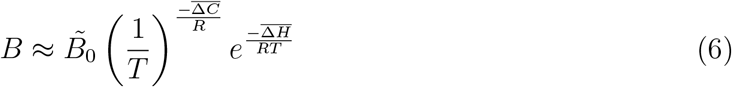

with the normalization constant, 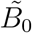, no longer depending on *h*. Note that the above correction for the enzyme level can also be applied to Eyring Eqs. (2) and (3), in which case they become mathematically identical to the Arrhenius relationship.
5. The micro and macro results, Eqs. (4) and (6), can be combined into a single expression for the temperature dependence of any variable, *Y* (*T*):

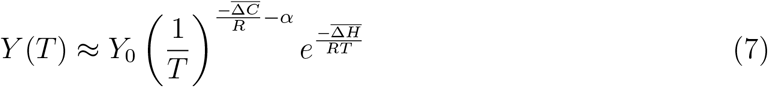

where *α* = 1 for the molecular level and 0 otherwise. *Y* (*T*) represents either a rate or various steady-state quantities [11] including variables that have been explicitly derived theoretically, such as in the MTE. For reaction rates at the molecular level *Y*_0_ is determined by Eq. (4). The corresponding extrema (either minima or maxima) in an Arrhenius plot now occur at 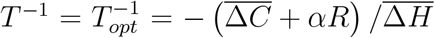. It should be noted that the thermodynamic parameters may have additional dependencies that make the forms of Eqs. (6) and (7) more complicated under certain conditions [11].

In addition to quantitatively explaining the origin and systematic curvature of the Arrhenius plot, our theory makes several further testable deductions that interrelate the key features of thermodynamic parameters (e.g. enthalpy and heat capacity), biological traits (e.g growth and metabolic rates), classic thermal traits (e.g. thermal range and optimum temperature). These various deductions, exhibited in Fig. S1, are summarized as follows:

i. The concave or convex form of the relationship between any biological trait and temperature (Eq. (4)-(7); Fig. 1).
ii. The relationship between differences in rates (e.g., *Y* (*T*_2_)*/Y* (*T*_1_)) and differences in temperatures (*T*_2_ − *T*_1_) (Eq. (S5.2)-(S5.3); Fig. S2).
iii. A linear relationship between heat capacity and enthalpy resulting from optimization of the rate (i.e. when the rate of change of *k* respect to temperature is zero), and where the slope of the relationship is the optimum temperature of the temperature response curve (Eq. (S6.2); Fig. S4).
iv. The linear relationship amongst all pairs of the key thermal traits of the temperature response curve such as the minimum, maximum, and optimum temperatures or thermal range (Eq. (S6.5.1-3); Fig. S6).
v. The linear relationships between a given thermal trait and fundamental thermodynamic parameters such as enthalpy (Eq. (S6.6.3-5); Fig. S7).
vi. The collapse, onto a single universal curve, of all temperature response curves after the appropriate re-scaling given by our theoretical framework (see discussion below and [11]; Eq. (9), (10); Fig. 2, Fig. S10). In particular our theory predicts that the optimum of this curve should be located at a rescaled temperature of 1.

**Figure 1.**
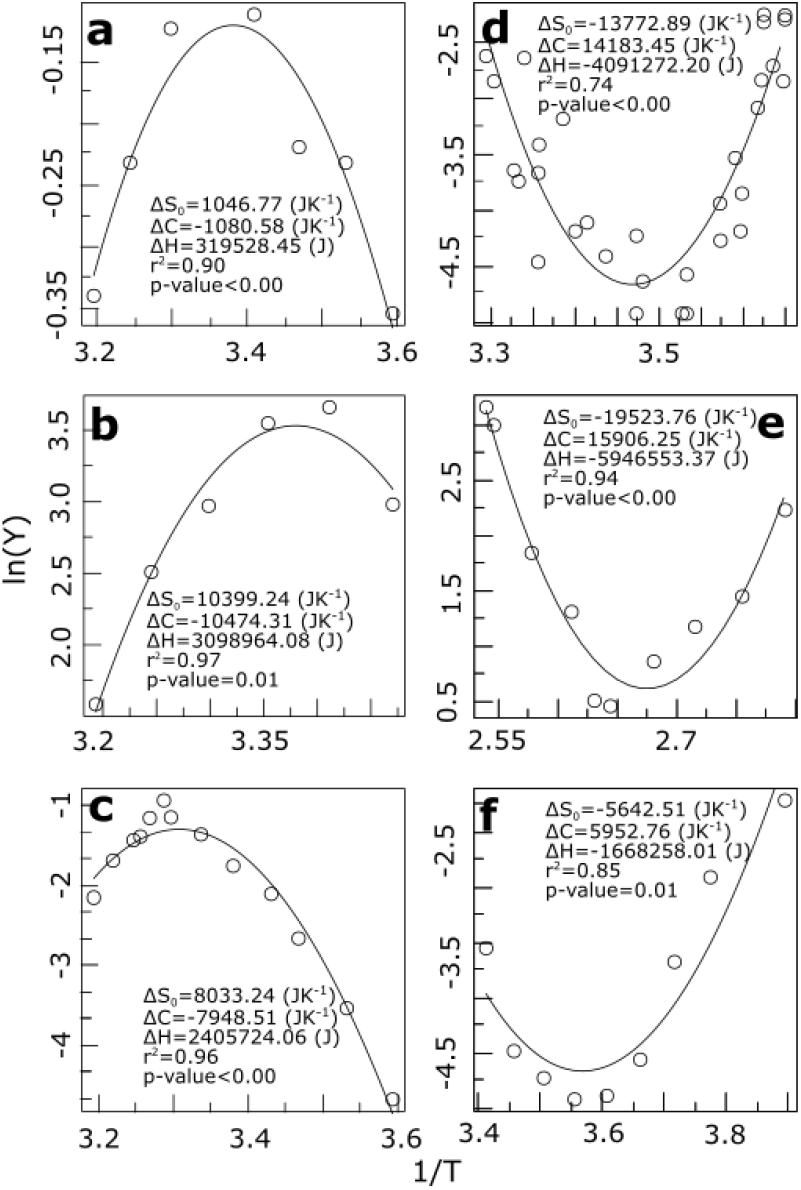
Temperature response curves compared to the predictions of Eqs. (5) and (7) for a wide diversity of biological examples. Plotted are ln(*Y*) vs. 1*/T* (in 1/K; where K is Kelvin degrees) showing (a)-(c) convex patterns and (d)-(f) concave patterns: (a) metabolic rate in the multicellular insect *Blatella germanica*, (b) maximum relative germination in alfalfa (for a conductivity of 32.1 dS/m), (c) growth rate in *Saccharomyces cerevisiae*, (d) mortality rate in the fruit fly (*Drosophila suzukii*), (e) generation time in strain 121, (f) metabolic rate in the rodent *Spermophilus parryii*. For references see Supplementary Methods. The x-axis is in units of (1*/K*) × 10^3^.

**Fig. 2.**
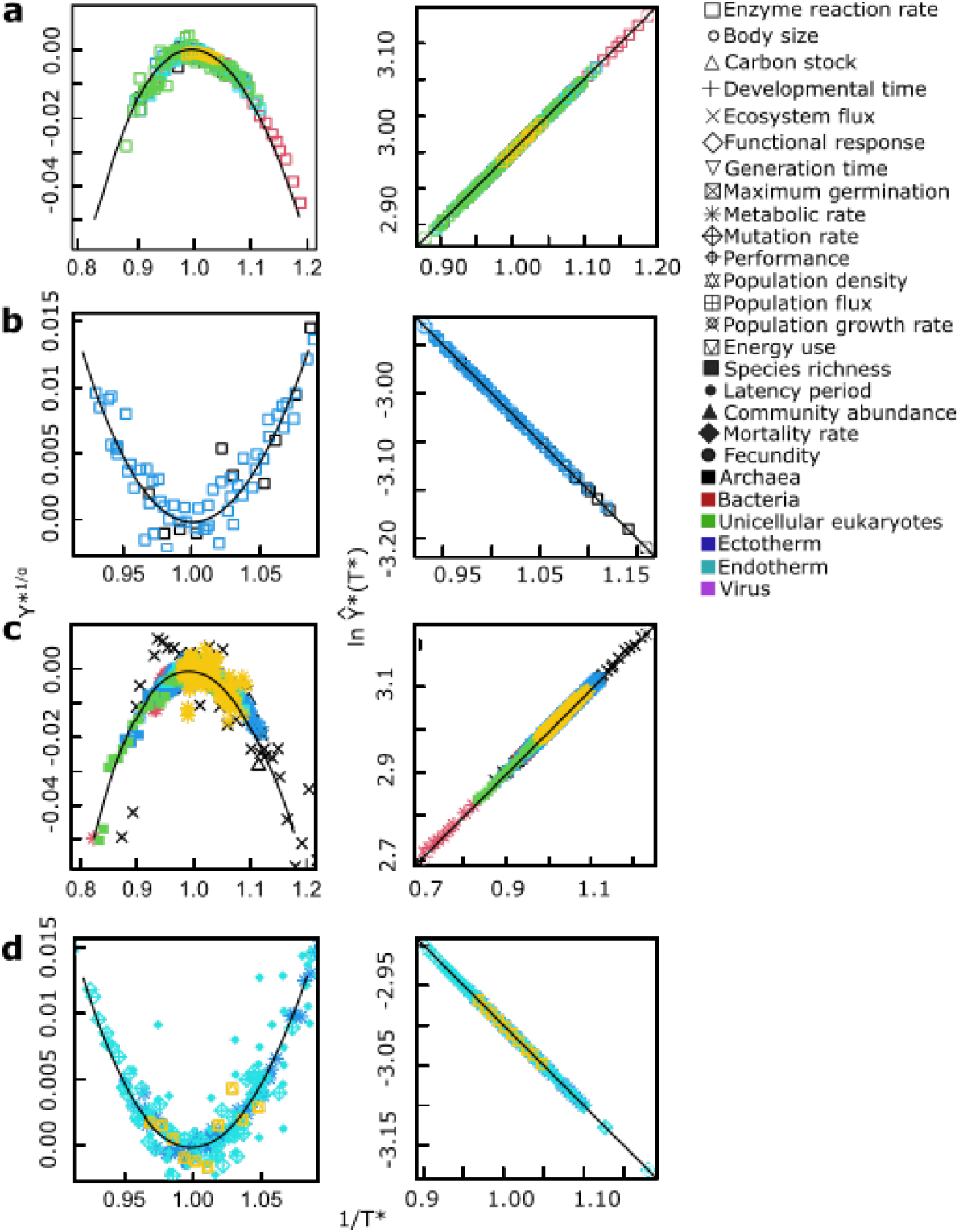
Universal patterns of temperature response predicted by Eqs. (9) and (10). The left panels show the convex and concave non-linear patterns predicted when ln *Y* ^*^ is plotted vs. 1*/T* ^*^, [Eq. (9)], whereas the right panels show the straight lines predicted when ln *Ŷ*^*^ is plotted vs. 1*/T* ^*^, [Eq. (10)]. All curves regardless of variable, environment and taxa collapse onto a single curve when plotted in either of these ways. These rescalings explicitly show the universal temperature-dependence of the data used in Fig. 1, as well as additional data from compiled studies. Panels (a) and (b) show molecular (enzymatic) data exhibiting the predicted concave and convex patterns on the left, while (c) and (d) show corresponding concave and convex patterns for data above the molecular level. Note that there appears to be no variance in the fits to the linear predictions (the right-hand set of graphs) whereas there is significant variation in the non-linear ones (the left-hand set of graphs). This is basically because ln(*Ŷ*^*^) >> ln(*Y* ^*^). The value of ln(*Y* ^*^) is typically around 0.01 with a variance much smaller than 0.005. Since ln(*Ŷ*^*^) = ln(*Y* ^*^) + *a* ln(*e/T* ^*^) and ln(*Ŷ*^*^) is typically around 3, fluctuations in ln(*Y* ^*^) are very much smaller and consequently completely lost. The point is that the difference between what is plotted in the left panels vs. that on the right, namely *a* ln(*e/T* ^*^), is in absolute value very large (more than 10 times the value of ln(*Ŷ*^*^); furthermore, it is almost a constant over the range of temperatures since it is logarithmic, whereas all of the temperature variation is in the much smaller term ln(*Ŷ*^*^).

Importantly, data fitting to deductions iii), iv), and v) all reveal universal relationships and constants. For example, the relationship between Δ*C* and Δ*H* holds across all data (fig S4) and is driven by a slope that is the optimum temperature associated to response curves.

### Comparing the theory to temperature response curve data across levels of biological organization and taxa

To assess the model performance, we compiled a database of 65 studies encompassing 128 temperature-response curves including those which are explicitly predicted by biological theories such as the MTE. Our survey included data of different rates/times/properties in different environments ranging from psychrophilic to hyperthermophilic organisms and across all domains of life, including viruses, bacteria, archaea and unicellular and multicellular eukaryotes covering both ectotherms and endotherms (see [11]).

We found that our theory provides an excellent fit to a wide variety of temperature response data for rates and times, spanning individual to ecosystem-level traits across viruses, unicellular prokaryotes, and mammals (see Supplementary Table S2). Fig. 1 shows some representative examples of fits to concave patterns with long tails at low and high temperatures (Fig. 1a-c) as well as convex patterns (such as the temperature dependence of endotherm metabolism and biological times, Fig. 1d-f) also with tails at both ends. Prediction ii) also fits the data well showing that curved temperature responses can be transformed into a linear relationship for discrete measures of both rates and temperatures (Fig. S2). As predicted in Eq. S6.2 we found a relationship between the estimated thermodynamic parameters − Δ*C* and Δ*H* (fig. S4) for all the (128) curve from our database.

Deductions iv-v) — the relationships among thermal traits and between thermal traits and parameters — are well supported by a subset of the overall data (Figs. S6 and S7).

### Universal scaling and data collapse

A powerful, but classic, method for exhibiting and testing the generality of a theory is to express it in terms of dimensionless variables which collapse the data onto a single “universal” curve across all scales [e.g. 30]. To do so here, we introduce dimensionless rates, *Y* ^*^, and temperatures, *T* ^*^, by rescaling them by *T*_*opt*_, where *Y* takes on either its minimum or maximum value, *Y*_*opt*_ = *Y* (*T*_*opt*_):

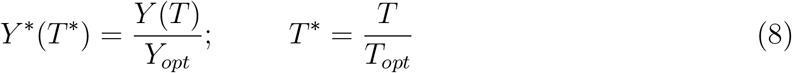

In terms of these rescaled variables, Eq. (7) reduces to the simple dimensionless form

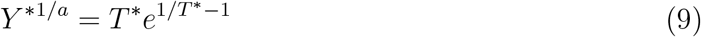

where 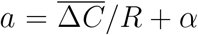 with *α* = 0 or 1, depending on whether the system is macro- or microscopic. Note that the optimum is given by 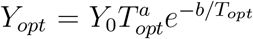 and *T_opt_* = −*b*/*a*, where 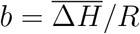 [11].

Our theory therefore predicts that when *Y* ^*1*/a*^ is plotted against 1*/T* ^*^ all of the various rates regardless of the specific processes collapse onto a single parameterless curve whose simple functional form is given by Eq. (9). Notice that this optimises at *T* ^*^ = 1 and encompasses in the same curve both the convex and concave behaviours predicted in the original Arrhenius plot as a function of *T*. In that regard, note also that the function

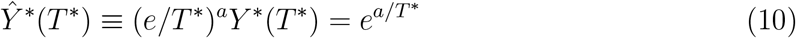

is predicted to be of a “pure” exponential Arrhenius form as a function of *T* ^*^. Thus, a plot of ln(*Ŷ*^*^(*T* ^*^)) vs. 1*/T* ^*^ should yield a straight line with slope *a* (see [11]).

Our prediction of the universal curve is very well supported by data, as illustrated in Fig. 2 where the collapse of all the data from this study for both convex and concave patterns regardless of organizational level, temperature range or taxa are shown. This result strongly supports the idea that our theory captures all of the meaningful dimensions of thermodynamic and temperature variation for diverse biological properties, which can ultimately be viewed as a single exponential relationship, Eq. (10). (See also Supplementary Material S8 and Fig. S8 for an alternative formulation for data collapse).

## Conclusion

In conclusion, we have derived a mechanistic yet simple theory for biological temperature responses. Our model is a general extension of the EEP equation, but unlike previous models requires only entropy to vary with temperature. From this single assumption, we derived six novel general predictions that include not only a formula for the temperature-dependence but also for explaining the parameters and relationships among thermodynamic properties, thermal traits, and between the two. This set of predictions leads to the discovery of universal constants, such an average global optimum for temperature response curves. We also derive a formula that expresses temperature dependence as a universal law that leads to data collapse across all levels of biological organization, taxa, and the whole range of temperature within which life can operate (−25 to 125°C). We do not imply that temperature is the only variable determining biological rates. We acknowledge the importance, and have included here, other variables that could be more limiting than temperature in certain environments, such as pH, which also determine enzymatic and other rates at higher levels of organization [31]. This framework allows us to make predictions for scenarios of global warming, disease spread, and industrial applications. Further extensions of this theory could incorporate time and other variables to predict the thermodynamic parameters or vice versa (i.e. the parameters could explain biological traits), and future connections could and should be made with non-equilibrium thermodynamics [32]. Finally, our framework allow us to better understand the diverse impacts of climate change upon processes at global scales, suggesting that processes such as mutation rates of viruses and mortality will likely increase, given their convex temperature response curves, but other such germination and growth rates will likely decrease given their concave temperature response curves (Fig. 1).

## Methods

Details on mathematical derivation, database compilation and estimation of parameters for the models are in Supplementary Methods.

## Data and code availability

The database and (R) code will be available in a public repository after acceptance. During the review process, data and code can be provided upon request.

## Acknowledgements

We thank authors that contributed with raw data and to Jim Brown for his comments on an early draft of this manuscript. JIA was supported by a Beca de Doctorado Nacional CONICYT 21130515. PAM was supported by grants AFB 17008 and ANID-FONDECYT 1200925 entitled “The emergence of of ecologies through metabolic cooperation and recursive organization.” JIA and GBW were supported by NSF 1838420, JIA and CPK were supported by NSF 1840301, GBW and CPK were supported by the Charities Aid Foundation of Canada (CAF) for the grant entitled “Toward Universal Theories of Ecological Scaling.” BD acknowledges support from projects ANID-FONDECYT 1150171 and 1190998.

## Author contributions

JIA and PM conceived the paper. JIA, CK, GW, and PM, derived the model. JIA compiled the database and made the statistical analysis. JIA, BD, CK, GW, PM wrote the paper.

## Competing interests

The authors declare no competing interests.

## Notes

### Competing Interest Statement

The authors have declared no competing interest.

### Summary of Updates

Details to improve on text, figures and tables

